# Mixtures of phthalates disrupt expression of genes related to lipid metabolism and peroxisome proliferator-activated receptor signaling in mouse granulosa cells

**DOI:** 10.1101/2024.05.02.592217

**Authors:** Hanin Alahmadi, Stephanie Martinez, Rivka Farrell, Rafiatou Bikienga, Nneka Arinzeh, Courtney Potts, Zhong Li, Genoa R. Warner

## Abstract

Phthalates are a class of known endocrine disrupting chemicals that are found in common everyday products. Several studies associate phthalate exposure with detrimental effects on ovarian functions, including growth and development of the follicle and production of steroid hormones. We hypothesized that dysregulation of the ovary by phthalates may be mediated by phthalate toxicity towards granulosa cells, a major cell type in ovarian follicles responsible for key steps of hormone production and nourishing the developing oocyte. To test the hypothesis that phthalates target granulosa cells, we harvested granulosa cells from adult CD-1 mouse ovaries and cultured them for 96 hours in vehicle control, a phthalate mixture, or a phthalate metabolite mixture (0.1-100 μg/mL). After culture, we measured metabolism of the phthalate mixture into monoester metabolites by the granulosa cells, finding that granulosa cells do not significantly contribute to ovarian metabolism of phthalates. Immunohistochemistry of phthalate metabolizing enzymes in whole ovaries confirmed that these enzymes are not strongly expressed in granulosa cells of antral follicles and that ovarian metabolism of phthalates likely occurs primarily in the stroma. RNA sequencing of treated granulosa cells identified 407 differentially expressed genes, with overrepresentation of genes from lipid metabolic processes, cholesterol metabolism, and peroxisome proliferator-activated receptor (PPAR) signaling pathways. Expression of significantly differentially expressed genes related to these pathways were confirmed using qPCR. Our results agree with previous findings that phthalates and phthalate metabolites have different effects on the ovary and interfere with PPAR signaling in granulosa cells.

## Introduction

Phthalates are a class of man-made chemicals that are widely known to be endocrine disruptors. They are commonly utilized in plastics and consumer products as plasticizers, solvents, and additives (Basso *et al*., 2022). Structurally, phthalates are composed of esters of ortho-phthalic acid with various hydrocarbon side chains (**Figure 1**). The varying side chains give phthalates a wide selection of properties and uses. Short-chain phthalates are most common in personal care products as they are used as fragrance excipients. Long-chain phthalates are commonly used as plasticizers for polyvinyl chloride (PVC) and are found in products such as vinyl flooring, medical tubing, and children’s toys (Panagiotou *et al*., 2021; Basso *et al*., 2022). As a result, humans are universally exposed to multiple phthalates as mixtures via dermal exposure from personal care products, oral exposure from food, and inhalation exposure from air and dust (Wang *et al*., 2019). Biomonitoring studies have found that women have greater exposure to phthalates than men, likely due to higher usage of personal care products, cosmetics, and cleaning products (Center for Disease Control and Prevention, 2018; Parlett *et al*., 2013). Furthermore, women from racial/ethnic minority groups have higher body burdens of phthalates than their white counterparts (Branch *et al*., 2015; Kobrosly *et al*., 2012). Phthalate exposure is associated with preterm birth, decreased fertility, and uterine and ovarian dysfunction (Radke *et al*., 2019; Basso *et al*., 2022; Hlisníková *et al*., 2020).

**Figure 1:**
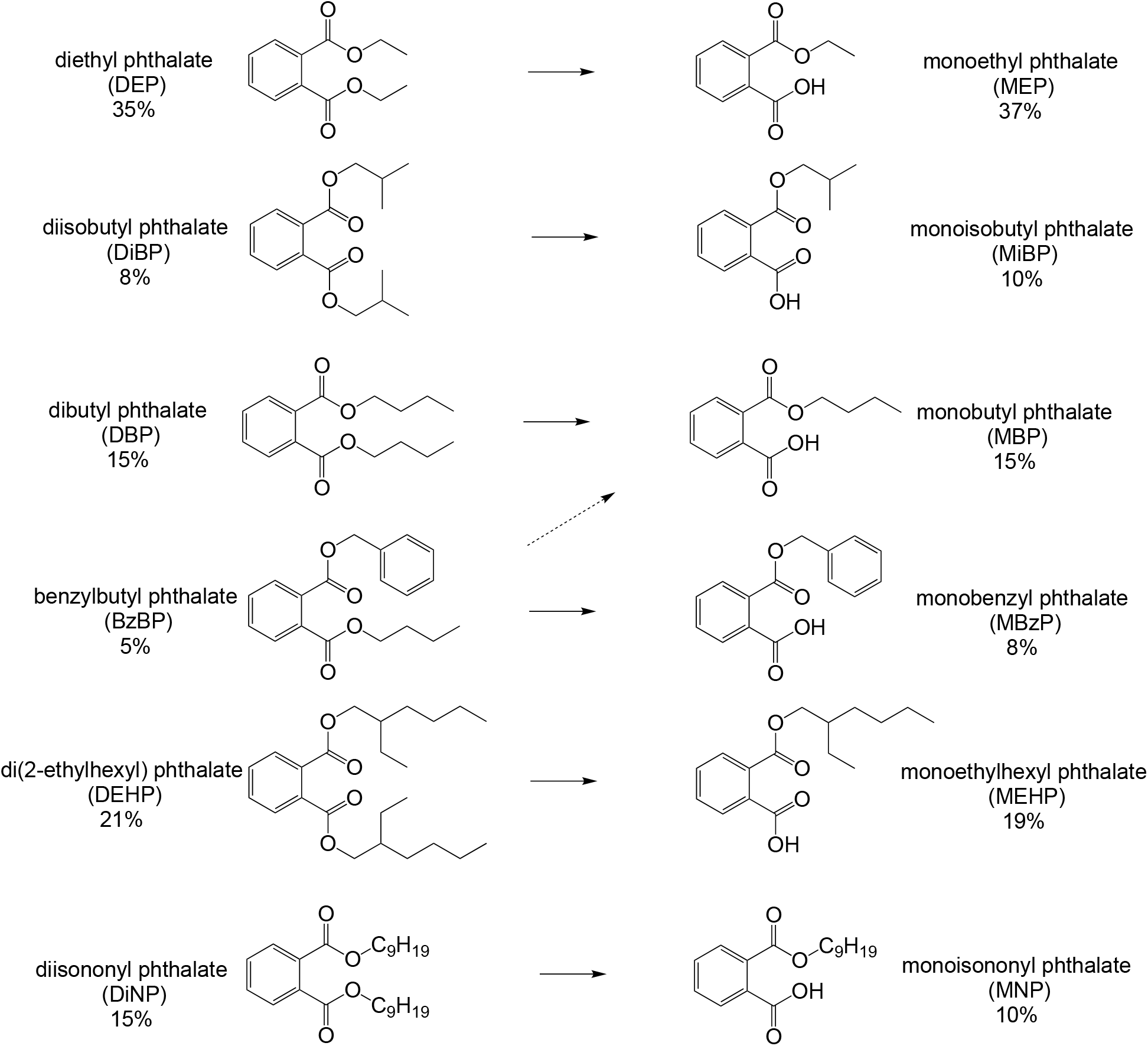
The phthalate mixtures used in this study are composed of diester phthalates (left) that correspond to the metabolites (right) measured in the urine of pregnant women in Illinois in the I-KIDS study. The differences in precents between diesters and their metabolites is due to the change in molecular weight of the compounds when one sides chain is hydrolyzed.

Although humans are exposed to diester phthalates in the environment, the diesters are easily converted to monoesters in the body. Numerous studies suggest that phthalate monoesters are the more potent toxicants (Lovekamp-Swan and Davis, 2003; Hannon *et al*., 2015; Wang *et al*., 2012). Initial metabolism is performed by lipases in the saliva and gut (Albro and Thomas, 1973). However, exposures that bypass oral metabolism can lead to higher concentrations of diester reaching other organs before further detoxification in the liver (Albro, 1986). Therefore, it is important to understand the capacity of target organs and the cells within them to bioactivate phthalates. Previously, we have found that mouse ovaries have a significant capacity to biotransform diester phthalates into monoesters (Warner *et al*., 2019). In addition, numerous laboratory studies have shown that phthalate mixtures of both diesters and metabolites have different effects on ovarian function compared to single phthalate exposures (Fletcher *et al*., 2022), emphasizing the need to study phthalate mixtures.

The ovary is composed of structure called follicles, which each contain and support the development of an oocyte. The most abundant populations of supporting cells in a follicle are the granulosa cells. As follicles grow and mature during the process of folliculogenesis, so do the granulosa cells, multiplying and differentiating to support the growth of the oocyte and participating in steroid hormone production. Previous studies have shown that phthalates have different effects across experimental models of life stages of the follicle (Warner *et al*., 2021; Zhou and Flaws, 2017), suggesting that phthalate toxicity varies at the tissue and/or cellular level due to diverse mechanisms of action and susceptibilities of certain cell types or stages of development. The key roles, abundance, and dynamic nature of granulosa cells across the stages of follicle development suggest that granulosa cells may be uniquely susceptible to disruption by phthalates. Indeed, a study of the impact of individual phthalate metabolites on cultured primary mouse granulosa cells showed that phthalate monoesters may act through endogenous peroxisome proliferator-activated receptor (PPAR) signaling (Meling *et al*., 2022). In cultured human granulosa cells, the same diester phthalate mixture used in this study was shown to disrupt progesterone signaling (Hannon *et al*., 2023).

Thus, herein we report on a study that contributes knowledge to our overall hypothesis that differences in metabolic capacity and phthalate toxicity among follicles at different stages of development are due to diversification in the cellular populations of follicles during folliculogenesis. We hypothesize that the different cell types express different levels of metabolizing enzymes and have different responses to phthalate exposure. In this paper, we focus on the granulosa cell, wherein we investigate whether granulosa cells are contributing to phthalate metabolism and the pathways through which granulosa cells are targets of phthalate toxicity using RNA-sequencing as a non-targeted approach.

## Methods

### Chemicals and Reagents

The phthalate mixture (PM) used in this study contains 35% diethyl phthalate (DEP), 21% di(2-ethylhexyl) phthalate (DEHP), 15% diisononyl phthalate (DiNP), 15% dibutyl phthalate (DBP), 8% diisobutyl phthalate (DiBP), and 5% benzylbutyl phthalate (BzBP) by weight (**Figure 1**). The phthalate metabolite mixture (MM) contains 37% monoethyl phthalate (MEP), 19% mono-(2-ethylhexyl) phthalate (MEHP), 15% monobutyl phthalate (MBP), 10% monoisononyl phthalate (MNP), 10% monoisobutyl phthalate (MiBP), and 8% mono-benzyl phthalate (MBzP) by weight (**Figure 1**). The compositions of the mixtures were derived from measured metabolites in the urine of pregnant women in Illinois (I-KIDS study, (Yazdy *et al*., 2018; Pacyga *et al*., 2022)). DEP (99.5%, catalog no. 524972), DBP (99%, catalog no. 524980), DiBP (99%, catalog no. 152641), BzBP (98%, catalog no. 308501), DEHP (99.5%, catalog no. D201154), and DiNP (99%, catalog no. 376663) were purchased from Sigma-Aldrich (St. Louis, MO). MEHP (catalog no. ARL-138S-CN) was purchased from AccuStandard (New Haven, CT). MEP (catalog no. M542580), MBP (catalog no. M525100), MiBP (catalog no. M54770), MBzP (catalog no. M524900), and MNP (catalog no. M567200) were purchased from Toronto Research Chemicals (North York, ON, Canada).

A stock solution of the pure PM was prepared as previously described (Zhou and Flaws, 2017) and stored at −20°C before use. As the metabolites are mostly powders at room temperature, a stock solution of 667 mg/mL of the MM in DMSO was prepared and stored at −20°C. Treatment solutions of 0.13, 1.33, 13.3, and 133 mg/mL were prepared by diluting the phthalate stocks in dimethyl sulfoxide (DMSO) such that the same volume of mixture and vehicle could be added per mL of culture media. The final working concentrations of phthalate mixtures in media were 0.100, 1.00, 10.0, and 100 µg/mL. Each working concentration contained the same volume of chemical and vehicle solution (0.75 µL of phthalate mixture in DMSO solution per mL of culture media equal to 0.075% DMSO).

The doses of the mixtures used in the culture experiments are representative of everyday, medical, and occupational levels of human exposure. Individual phthalates within the mixture doses of 0.1 and 1.0 µg/mL are similar to everyday exposure measured in urine in the National Health and Nutrition Examination Survey (NHANES) in the ng/mL range and follicular fluid levels measured in *in vitro* fertilization patients (Center for Disease Control and Prevention, 2018; Du *et al*., 2016; Krotz *et al*., 2012). Individual phthalates within the mixture doses of 10 and 100 µg/mL encompass medical and occupational exposure to phthalates as measured in urine, serum, and blood (Hauser *et al*., 2004; Hernández-Díaz *et al*., 2009; Hines *et al*., 2012; Sjoberg *et al*., 1985).

### Animals

Adult CD-1 mice were purchased from Charles River Laboratories (Wilmington, MA) and allowed to acclimate to the facility before use. The mice were housed at the University of Illinois at Urbana-Champaign, Veterinary Medicine Animal Facility. Food (Harlan Teklad 8604) and water were provided for ad libitum consumption. Temperature was maintained at 22 ± 1 °C, and animals were subjected to 12-h light-dark cycles. The Institutional Animal Use and Care Committee at the University of Illinois at Urbana-Champaign approved all procedures involving animal care, euthanasia, and tissue collection.

### Granulosa cell culture

Isolated granulosa cells from antral follicles were cultured in the presence of vehicle control (DMSO), phthalate mixture or metabolite mixture (0.1, 1, 10, or 100 µg/mL, n = 4–9 replicates) as previously described (Meling *et al*., 2022). Briefly, granulosa cells were harvested from the ovaries of 5– 10 32–42 day old CD-1 mice by tearing the tissue apart with watchmaker forceps in media composed of α-minimal essential media (α-MEM, Life Technologies, Grand Island, NY) with 1 × ITS (10 μg/mL insulin, 5.5 μg/mL transferrin, 5ng/mL sodium selenite, Sigma-Aldrich, St. Louis, MO), 1 × antibiotic antimycotic (100 U/mL penicillin, 100 μg/mL streptomycin, 25ng/mL amphotericin B (Sigma-Aldrich), 10% FBS (Atlanta Biologicals), and 5 IU/mL follicle stimulating hormone (Dr. A.F. Parlow). Cell suspensions were passed through a 23 gauge needle and 40 μm strainer and then expanded in culture for 3–4 days at 37 °C and 5% CO_2_. Cells were released with 0.25% trypsin-EDTA and replated at 1 × 10^5^ cells/well in 12-well plates. The next day, media were exchanged of phthalate-supplements treatments, and were not changed over the 96 hour culture period. Additional control wells with media only (no cells) were kept in the incubator for the duration of the experiment to analyze ambient degradation of phthalates. At the end of the culture period, media were collected for mass spectrometry analysis of phthalate content to analyze metabolism and enzyme linked immunosorbent assays (ELISAs) to measure hormone levels. Cells were flash frozen and stored at −80 °C until RNA extraction for qPCR and bulk RNA sequencing analysis.

### Phthalate metabolism analysis

Media from 96 hour granulosa cell cultures were subjected to mass spectrometry analysis of phthalate parent compounds and metabolites as previously described (Warner *et al*., 2019). All experiments contained appropriate control wells with no tissue to account for ambient degradation of phthalates and phthalate free controls to detect contamination from the lab environment. No phthalates were detected in phthalate free controls. All six parent compounds were measured as well as their respective primary monoester metabolites MEP, MBP, MiBP, MBzP, MEHP, and MiNP. Media samples were stored at −80°C until analysis and were injected directly into the instrument. Samples were analyzed with the 5500 QTRAP LC/MS/MS system (Sciex, Framingham, MA) in the Metabolomics Lab of the Roy J. Carver Biotechnology Center, University of Illinois at Urbana-Champaign. Software Analyst 1.6.2 was used for data acquisition and analysis. The 1200 series HPLC system (Agilent Technologies, Santa Clara, CA) includes a degasser, an autosampler, and a binary pump. The LC separation was performed on a Phenomenex (Torrance, CA) Gemini C6-phenyl column (2 x 100 mm, 3 μm) with mobile phase A (0.1 % formic acid in water) and mobile phase B (0.1 % formic acid in acetontrile). The flow rate was 0.25 mL/min. The linear gradient was as follows: 0-1 min, 70 % A; 10–15 min, 50 % A; 20–27.5 min, 10 % A; 28–34 min, 70 % A. The autosampler was set at 10°C. The injection volume was 5 μL. Mass spectra were acquired under both positive electrospray ionization (ESI) with the ion spray voltage of 5000 V and negative ESI with the ion spray voltage of −4500 V. The source temperature was 500°C. The curtain gas, ion source gas 1, and ion source gas 2 were 32, 50, and 65 psi, respectively. Multiple reaction monitoring was used for quantitation. Percent metabolism of each compound during the culture period was calculated to account for ambient degradation of phthalates and represents the percent decrease in actual phthalate concentration in the media from the beginning of the experiment until the end. Percent metabolism is not compared between the culture systems because of the different amounts of tissue in culture and the different experimental lengths. DiNP metabolism was not able to be reported because DiNP is a complex mixture of isomers, which makes targeted analysis of metabolites tricky (Saravanabhavan and Murray, 2012). We selected a monoester metabolite that we expected to be a major isomer (MNP, **Figure 1**), but it was not detected in any samples. We do not report metabolism from the 0.1 μg/mL treatment group because of contamination identified by the mass spectrometry analysis in the vehicle control group at similar concentrations as the phthalate measurements in the 0.1 μg/mL PM treatment group for some phthalates.

### Immunohistochemistry staining for metabolizing enzymes

Whole, untreated and uncultured ovaries from 8 day old and 32 – 42 day old CD-1 mice were fixed in 4% paraformaldehyde overnight and transferred to 70% ethanol for immunohistochemical analysis. Ovaries were embedded in paraffin wax and serial sectioned at 5 µm. Between two and six sections spanning the length of the ovary were mounted on a glass slide. Slides were randomly assorted to be stained in multiple batches. Following deparaffinization, the tissues were subjected to heat-induced antigen retrieval (10 mM sodium citrate butter at pH 6.0) for 15 minutes in a pressure cooker and then blocked in 3% hydrogen peroxide for 10 minutes. Slides were then incubated with 2.5% normal horse serum from the ImmPRESS Horse Anti-Rabbit IgG Polymer Kit (catalog no. MP-7401, Vector Laboratories) for 20 minutes, rabbit polyclonal primary antibody for 60 minutes (ab52492, Abcam and sc-373759, Santa Cruz), and ImmPRESS Horse Anti-Rabbit IgG Polymer Reagent secondary antibody (catalog no. MP-7401, Vector Laboratories) for 30 minutes. ImmPACT Diaminobenzidine (DAB) peroxidase substrate solution (SK-4105, Vector Laboratories) was then applied until color optimally developed. Each sample was exposed to the chromogen for equal amounts of time. The slides were then rinsed, counterstained with Tasha’s Automated Hematoxylin, and cover-slipped. Primary and secondary negative controls were subjected to the same methods listed above, except instead of antibody, the negative control tissues were incubated with tris buffered saline. Negative controls were confirmed to have no positive staining after exposure to the chromogen for equal amounts of time as the samples. Images were captured using a NanoZoomer Digital Pathology System (Hamamatsu) and analyzed using the FIJI version of ImageJ (https://imagej.net/Fiji).

### Hormone measurements

Culture media were collected after 96 hours of granulosa cell culture from 4–9 separate cultures and subjected to enzyme-linked immunosorbent assays (ELISAs, DRG International Inc., Springfield, New Jersey) for measurement of pregnenolone and progesterone. Assays were run according to the manufacturer’s instructions. Samples were run in duplicate and diluted if necessary (to match the dynamic range of each ELISA kit). Progesterone coefficients of variability were 9% (intraassay) and 12% (interassay). Pregnenolone coefficients of variability were at 6% (intraassay) and 10% (interassay).

### Bulk RNA sequencing analysis

Total RNA was isolated from snap-frozen granulosa cells using a MicroElute Total RNA Kit (Omega Bio-Tek, Norcross, GA) according to the manufacturer’s instructions. RNA was eluted in RNase-free water and the concentration was determined using a NanoDrop (λ = 260nm, Nanodrop Technologies, Inc., Wilmington, DE). Five samples each from the DMSO, 0.1 μg/mL PM, 10 μg/mL PM, 0.1 μg/mL MM, and 10 μg/mL MM treatment groups were selected for RNA sequencing analysis. Sequencing was performed at the Roy J. Carver Biotechnology Center at the University of Illinois at Urbana-Champaign. The RNAseq libraries were prepared with Illumina’s TruSeq Stranded mRNAseq Sample Prep kit (Illumina). Reads were 100 nt in length with Read 1 aligned to the ANTISENSE strand. The libraries were pooled and quantitated by qPCR and sequenced on one SP lane for 101 cycles from one end of the fragments on a NovaSeq 6000. Fastq files were generated and demultiplexed with the bcl2fastq v2.20 Conversion Software (Illumina). Adaptors were trimmed from the 3’-end of the reads. Reads were then checked for quality using FASTQC (version 0.11.8) on individual samples then was summarized into a single html report by using MultiQC version 1.9 (Ewels *et al*., 2016). Average per-base read quality scores were over 30 in all samples, so no additional trimming was performed.

All analyses from summation of counts to the gene level (including the codes in .Rmd file) were done on a laptop in R version 4.0.4 using packages as indicated below. Reads were aligned to NCBI’s *Mus musculus* GRCm39 genome. Salmon version 1.2.1 (Patro *et al*., 2017) was used to quasi-map reads to the transcriptome and quantify the abundance of each transcript. The transcriptome was first indexed using the decoy-aware method in Salmon with the entire genome file as the decoy sequence. Then quasi-mapping was performed to map reads to transcriptome with additional arguments to correct sequence-specific and GC content biases, compute bootstrap transcript abundance estimates and help improve the accuracy of mappings. Gene-level counts were then estimated based on transcript-level counts using the “bias corrected counts without an offset” method from the tximport package. This method provides more accurate gene-level counts estimates and keeps multi-mapped reads in the analysis compared to traditional alignment-based method (Soneson *et al*., 2015). Percentage of reads mapped to the transcriptome ranged from 84.2 to 89.2%. The unmapped reads were discarded and the number of remaining reads (range: 16.6 - 24.1 million per sample) were kept for further analysis. Trimmed mean of M values (TMM) normalization (Robinson and Oshlack, 2010) was performed in the edgeR package (Robinson *et al*., 2010). The detection threshold was then set at 1 count per million (cpm) in at least 4 samples, which resulted in 25,769 genes being filtered out, leaving 13,484 genes to be analyzed for differential expression that contain 99.90% of the reads. After filtering, TMM normalization was performed again and normalized log2-based count per million values (logCPM) were calculated using edgeR’s (McCarthy *et al*., 2012) cpm() function with prior.count = 2 to help stabilize fold-changes of extremely low expression genes. Multidimensional scaling in the limma (Ritchie *et al*., 2015) package was used as a sample quality control step to check for outliers or batch effects. The normalized logCPM values of the top 5,000 most variable genes were chosen to construct the multidimensional scaling plot. Observed batch effects were accounted for by adding batch groups as coefficients on the statistical model run on the normalized logCPM values. Differential gene expression analysis was performed using the limma-trend method (Chen *et al*., 2016) using a model of Group + Batch. Pairwise comparisons of each treatment compared to DMSO were made and also combined in one F-test to calculate a oneway ANOVA across the 5 treatments. Multiple testing correction was done using the False Discovery Rate (FDR) method (Benjamini and Hochberg, 1995).

Data obtained from RNA sequencing were functionally analyzed using The Database of Annotation, Visualization, and Integrated Discovery Bioinformatics (DAVID) 2021 following the previously published protocol (Huang *et al*., 2009b, 2009a). Genes with an ANOVA FDR p-value < 0.05 were entered into DAVID for functional annotation analysis for a total of 407 genes. “Gene_Ontology” and “Pathways” and the denoted DAVID defined defaults were selected for functional annotation clustering. To determine if functional gene groups were valuable, annotation clusters with a significant enrichment score ≥ 1 were further explored (Huang *et al*., 2009b) using epigenetic Landscape In Silico deletion Analysis (LISA) tool at cistrome.org (Qin *et al*., 2020) and the ppargene.org database.

### Gene expression analysis

Total RNA was isolated from snap-frozen granulosa cells using a MicroElute Total RNA Kit (Omega Bio-Tek, Norcross, GA) according to the manufacturer’s instructions. RNA was eluted in RNase-free water and the concentration was determined using a NanoDrop (λ = 260nm, Nanodrop Technologies, Inc., Wilmington, DE). RNA (400 ng) was reverse-transcribed to complementary DNA (cDNA) using iScript Reverse Transcriptase (Bio-Rad Laboratories, Inc., Hercules, CA) according to the manufacturer’s protocol. Quantitative polymerase chain reaction (qPCR) analysis was done in 96-well plates in 10 μL reaction volumes containing 0.75 pmol/μL for each of two primers, cDNA derived from 6.67 ng total RNA, and 1 × SsoFast EvaGreen dye (Bio-Rad Laboratories). Quantifications were done using the CFX Duet Real-Time Detection System (Bio-Rad Laboratories) and CFX Manager Software. All qPCR primers (Integrated DNA Technologies, Coralville, IA) were purchased lyophilized and dissolved in DNase-free water and are listed in **Table S1**. A mean value of duplicate runs was calculated for all samples. The qPCR protocol began with incubation at 95 °C for 5min. This was followed by 40 cycles at 95 °C for 10s, at 60 °C for 10s, and at 72 °C for 10s. Melting from 65 °C to 95 °C was followed by extension at 72 °C for 2min. Melting temperature graphs, standard curves, and threshold cycle (Ct) values were acquired for each gene analyzed. The expression data were normalized to the corresponding values for Beta Actin (*ActB*). Individual relative fold changes compared to the vehicle control treatment group were calculated using the Pfaffl method (Pfaffl, 2001).

### Statistical Analysis

Data were expressed as the mean ± standard error of the mean (SEM). Data were analyzed by comparing treatment groups to control using IBM SPSS version 26 software (SPSS Inc., Chicago, IL, USA). Outliers were removed by the Grubb’s test using GraphPad outlier calculator software (GraphPad Software Inc., La Jolla, CA). All data were continuous and assessed for normal distribution by Shapiro-Wilk analysis. If data met assumptions of normal distribution and homogeneity of variance, data were analyzed by one-way analysis of variance (ANOVA) followed by Tukey HSD or Dunnett 2-sided *post-hoc* comparisons. However, if data met assumptions of normal distributions, but not homogeneity of variance, data were analyzed by ANOVA followed by Games-Howell or Dunnett’s T3 *post-hoc* comparisons. If data were not normally distributed or presented as percentages, the independent sample Kruskal-Wallis H followed by Mann-Whitney U non-parametric tests were performed. For all comparisons, statistical significance was determined by p-value ≤ 0.05. If p-values were greater than 0.05, but less than 0.10, data were considered to exhibit a trend towards significance.

## Results

### Metabolism of phthalates by cultured granulosa cells

The ability of granulosa cells to metabolize phthalates *in vitro* was assessed by comparing the concentration of parent phthalates in the treatment at day 0 of culture compared to the concentration of metabolites present at the end of the 96-hour culture period. The concentration of metabolites over the same time period in media without cells was subtracted from this value to account for non-biological degradation of phthalates. Results are reported as the percent metabolism of each individual parent phthalate in the mixture (**Figure 2**). Granulosa cells metabolized the largest percent of DiBP, DBP, and BzBP, which is consistent with our previous findings for antral follicles. Phthalate metabolism topped out at 15-20% of each phthalate, meaning that we detected no more than 20% of the expected concentration of each metabolite given the initial dose. Only 5% of the phthalates at 100 ug/mL were metabolized, suggesting that higher concentrations of phthalates overwhelm the ability of enzymes to clear them. On average 105% of the initial concentration of phthalates in the control wells, with no cells, were recovered at the end of the culture period, suggesting slight environmental contamination in the laboratory or from the mass spectrometer. For the 10 and 100 ug/mL doses in the wells with granulosa cells, total phthalate recovery was only 52% (range: 41 – 74%), suggesting that phthalates, especially at the highest does, are migrating into the cells.

**Figure 2:**
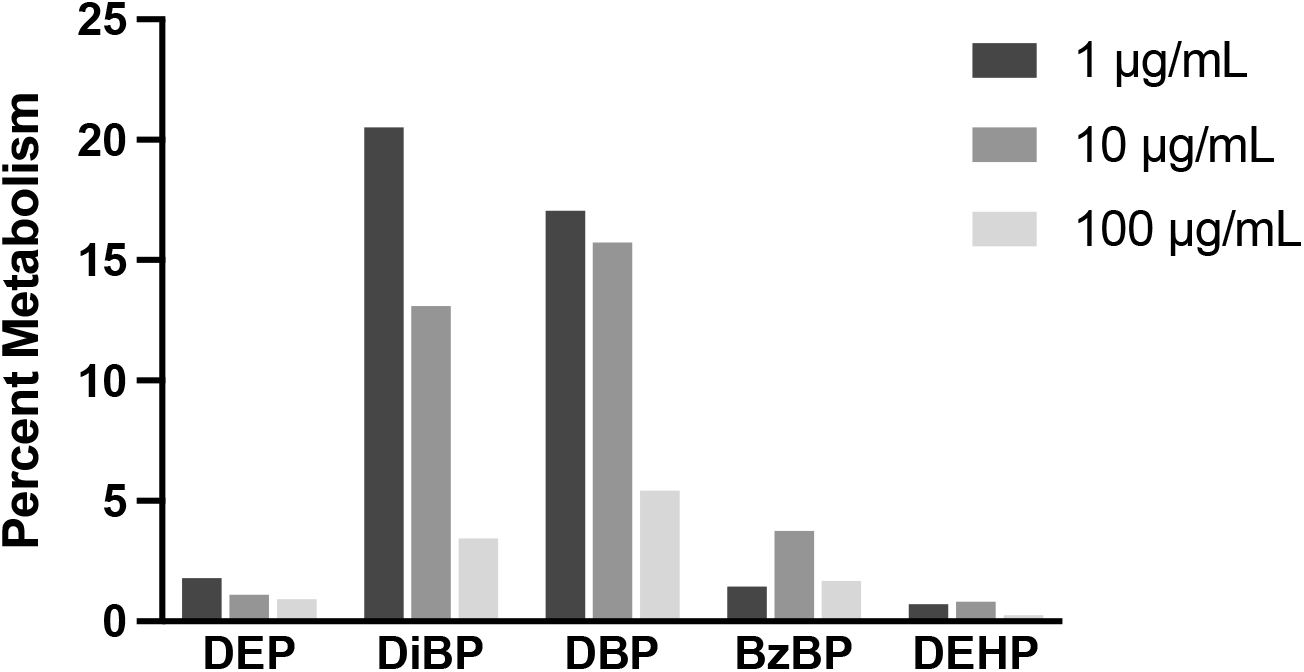
Percent metabolism of individual phthalates measured by LC-MS/MS following culture of granulosa cell from adult mouse antral follicles in the presence of the phthalate mixture (1–100 μg/mL) for 96 hrs.

### Role of metabolizing enzyme expression in ovarian cell types

Previously, we identified the phthalate-metabolizing enzymes lipoprotein lipase (LPL) and aldehyde dehydrogenase family 1, subfamily A1 (ALDH1A1) in antral follicles and whole prenatal mouse ovaries via western blot and confirmed expression of *Lpl* and *Aldh1a1* via qPCR (Warner *et al*., 2019). We identified the highest levels of LPL in neonatal ovaries and ALDH1A1 in antral follicles. Neonatal ovaries contain predominantly primary and primordial follicles with different populations of cells compared to the more developed antral follicles, leading us to hypothesize that expression of these enzymes varies across cell types and across different stages of development in the ovary. Relative gene expression and protein levels of these enzymes concurred between our previous analyses, but cell-specific information was not available. To spatially identify enzyme levels across cell types at different stages of follicle development, we performed immunohistochemistry staining in whole, untreated ovaries from 8-day-old neonatal mice and young adult mice.

In neonatal ovaries, little to no expression of ALDH1A1 was observed (**Figure 3A**), whereas LPL was detected in the early cuboidal granulosa cells in primary follicles and in the stroma between follicles (**Figure 3B**). This is consistent with our previous finding that LPL was more strongly expressed than ALDH1A1 in neonatal ovaries.

**Figure 3:**
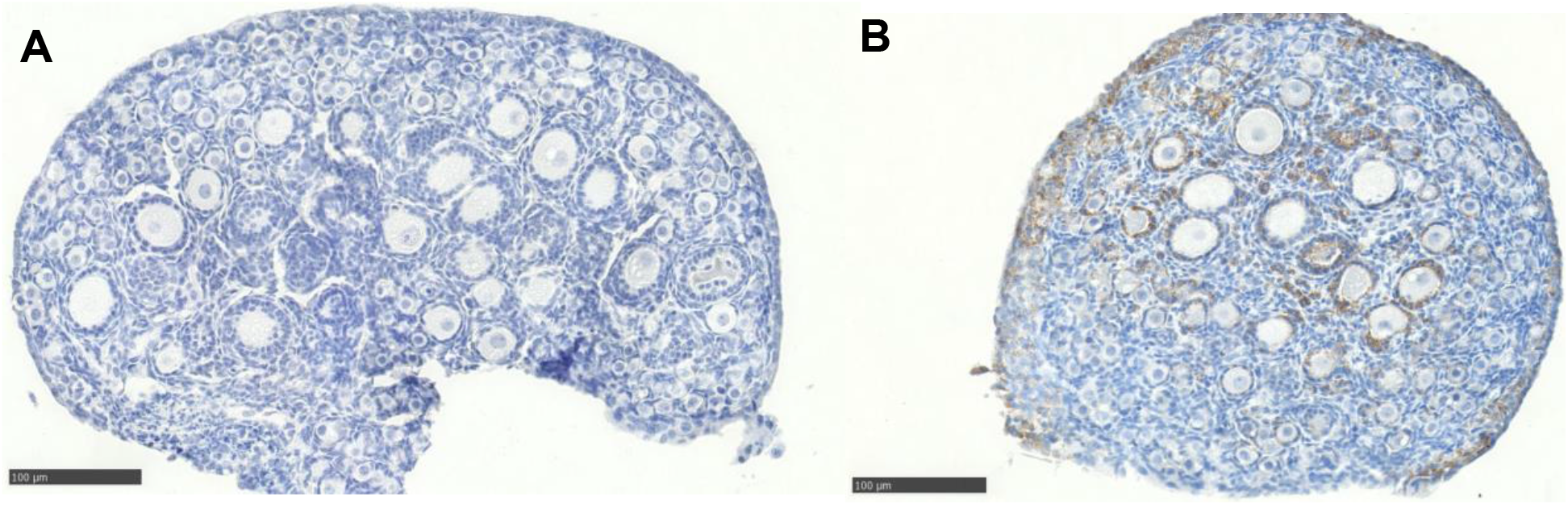
Immunohistochemical staining of whole, uncultured, untreated ovaries from 8-day old neonatal mice. (A) Representative image of ALDH1A1 staining, which was not observed. (B) Representative image of LPL staining shows staining (brown) in the granulosa cells of primary follicles and in the stroma.

Interestingly, in adult ovaries, the stroma surrounding the follicles exhibited the strongest ALDH1A1 staining (**Figure 4A-D**). Primary, preantral, and antral follicles did not express ALDH1A1, except for the theca cells of antral follicles (**Figure 4C-D**, arrows). Like the neonatal ovaries, positive staining was observed for LPL in the stroma and granulosa cells of primary follicles as well as preantral follicles (**Figure 4E-H**). However, granulosa cells of antral follicles did not stain for LPL, suggesting that expression of this enzyme decreases as folliculogenesis progresses. Minimal expression of phthalate-metabolizing enzymes in granulosa cells from antral follicles is consistent with our observation of low rates of biotransformation of phthalates in granulosa cell cultures.

**Figure 4:**
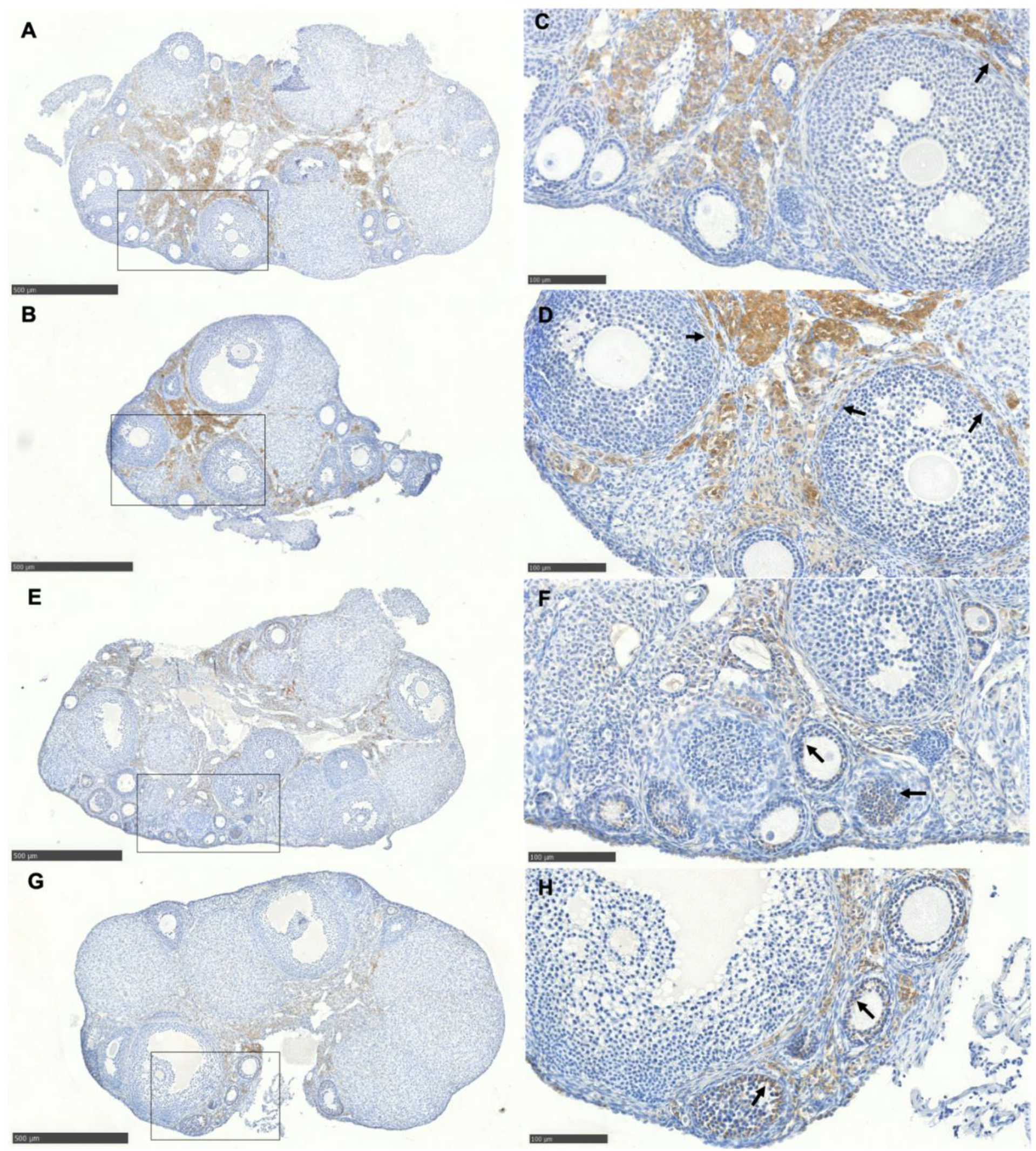
Immunohistochemical staining of uncultured, untreated young adult mouse ovaries. Images on the right are insets of the black boxes on the right. (A-D) Representative images of ALDH1A1 staining (brown) in the stroma and theca cells of antral follicles (arrows) (E-H) Representative images of LPL staining (brown) show staining in the stroma and granulosa cells of pre-antral follicles (arrows).

### Effects of the phthalate mixtures on steroidogenesis in granulosa cells

Although theca cells are required for the full steroidogenic pathway in the synthesis of estradiol, granulosa cells are capable of producing progesterone on their own. The steroidogenic acute regulatory protein (STAR) imports cholesterol into the ovary. Cholesterol is converted into pregnenolone by CYP11A1, followed by conversion into progesterone by HSD3B1. Granulosa cells are also essential for performing the final conversion of estrone into estradiol via HSD17B1. To investigate if the phthalate treatments altered the capacity of granulosa cells to these initial steps in the steroidogenic processes, we measured concentrations of pregnenolone and progesterone in media from the granulosa cell cultures and expression of *Star*, *Cyp11a1,* and *Hsd3b1* in extracted RNA from the granulosa cells. The concentrations of pregnenolone and progesterone were not altered compared to control in any treatment group (**Figure 5A**). However, the expression of *Cyp11a1* and *Hsd3b1* were statistically significantly decreased in the 100 μg/mL PM treatment groups compared to control, whereas expression of *Star* was significantly increased in the 1 and 100 μg/mL MM groups and decreased in the 10 and 100 μg/mL MM groups compared to controls (**Figure 5B**). We also measured expression of *Hsd17b1* and found that 10 and 100 μg/mL of the PM significantly decreased expression compared to control.

**Figure 5:**
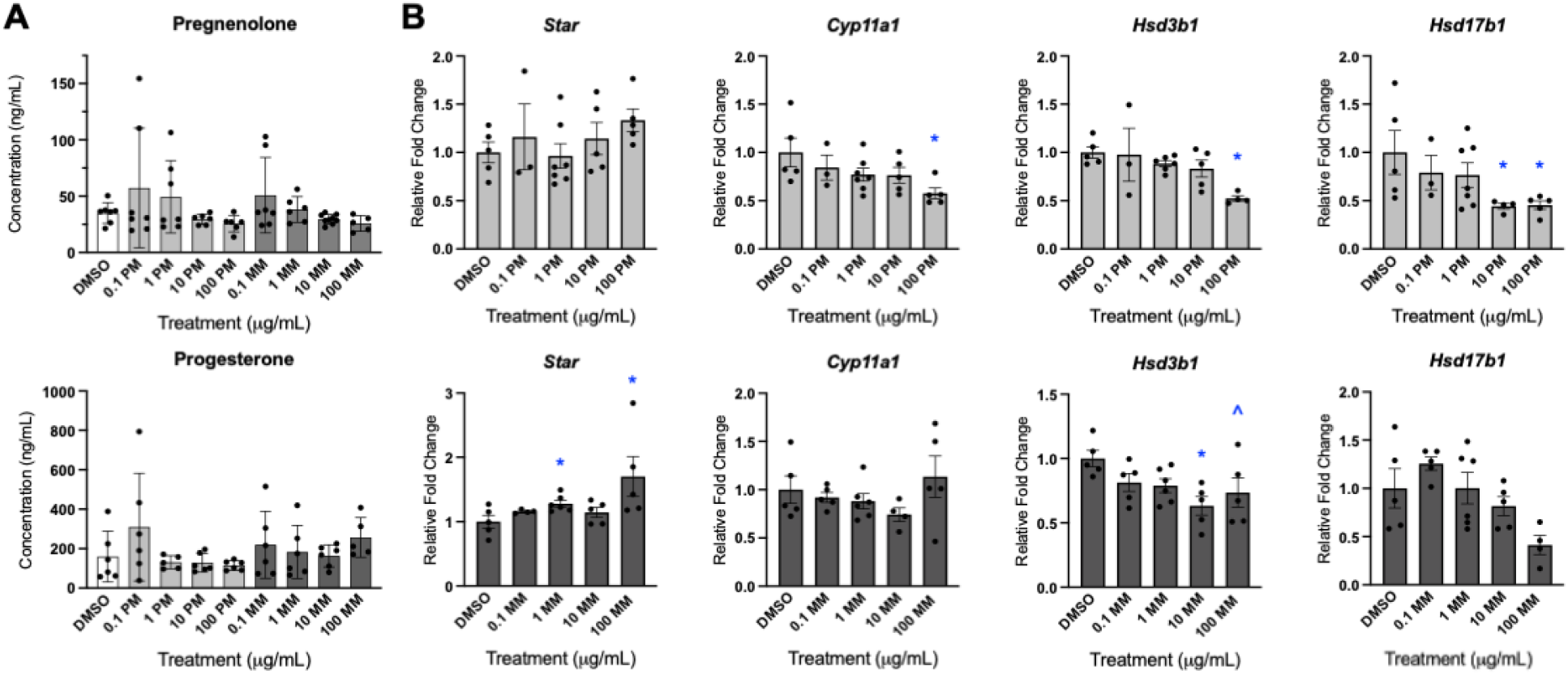
Impact of phthalate exposure on steroidogenesis in granulosa cells. Granulosa cells were harvested from adult female mice and cultured for 96 hours in the presence of the phthalate mixture (PM) or metabolite mixtures (MM) at 0.1 – 100 μg/mL. (A) Pregnenolone and progesterone concentrations in culture media collected at the end of the culture period (B) Gene expression measured via qPCR of key steroidogenesis enzymes in granulosa cells collected at the end of the culture period. Graphs represent mean ± SEM from 3–6 separate experiments. Asterisks (∗) indicate significant differences from the control (p ≤ 0.05) and ^ indicates a trend toward significance (p ≤ 0.10).

### Gene expression analysis of the effect of phthalate mixtures on granulosa cells

Extracted RNA from granulosa cells cultured with DMSO or 0.1 or 10 μg/mL of PM or MM were subjected to bulk RNA-seq analysis to investigate the differences in gene expression in response to phthalate exposure. With a false discovery rate < 0.2, 1438 differentially expressed genes were identified (**Figure 6A**), with the 10 μg/mL PM treatment group showing the most differentially expressed genes compared to the control (**Figure 6B**). The FDR p-value was restricted to < 0.05 for further analysis. Using a one-way ANOVA across all 5 treatment groups yielded 407 genes that showed a difference somewhere across the treatments (**Table S2**). With the restricted FDR, the 0.1 μg/mL PM treatment group had one upregulated gene, the 10 μg/mL PM treatment group has 66 downregulated genes and 57 upregulated genes, the 0.1 μg/mL MM treatment group had 2 downregulated genes and 13 upregulated genes, and the 10 μg/mL MM treatment group had 29 downregulated genes and 61 upregulated genes (**Figure 6C**). Twenty-one genes were altered in more than one treatment group (**Figure 6D**).

**Figure 6:**
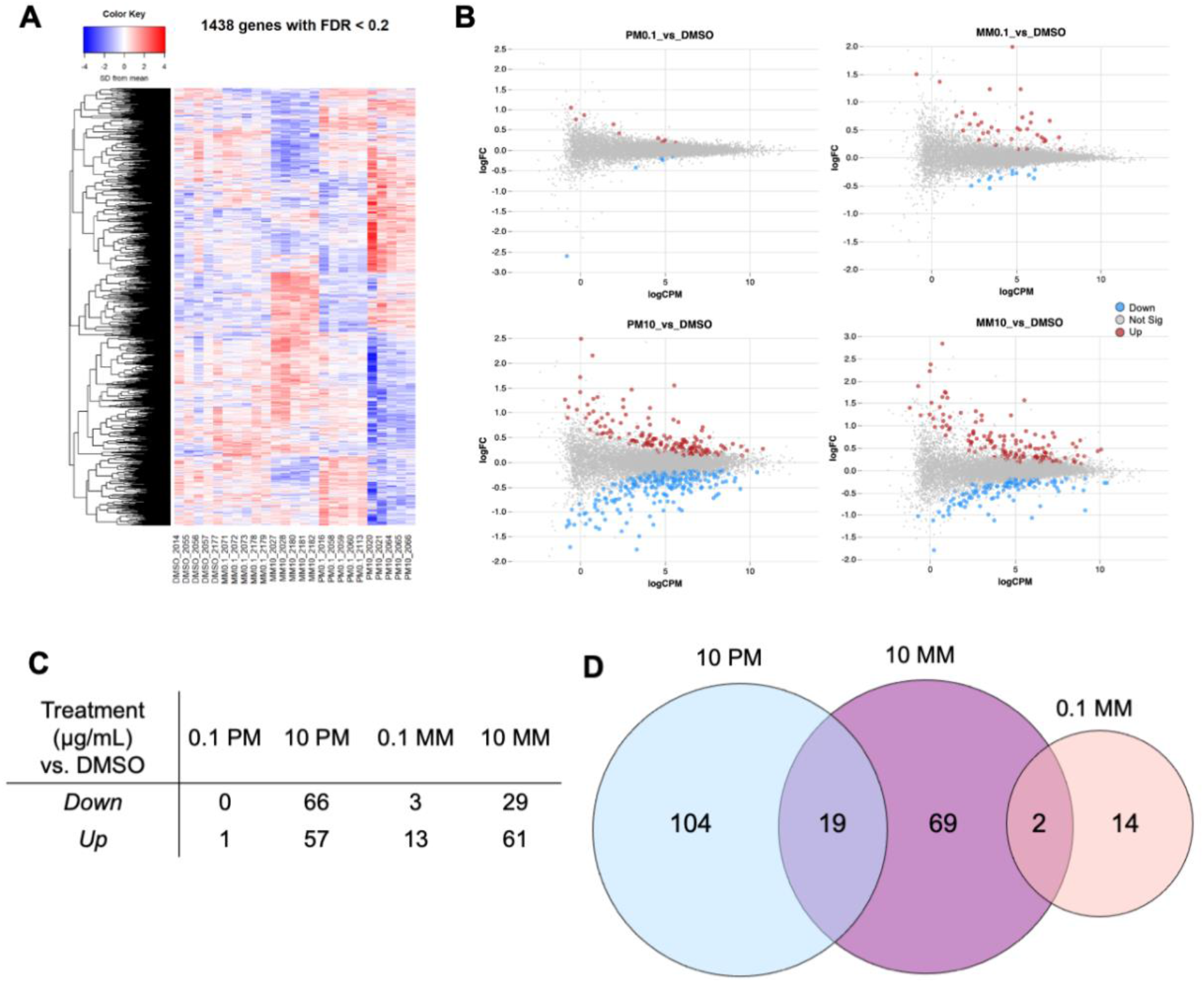
RNA-sequencing of granulosa cells exposed to the phthalate mixtures. Granulosa cells harvested from antral follicle from young adult mouse ovaries were cultured in the presence of DMSO (vehicle control), 0.1 μg/mL of the phthalate mixture (PM0.1), 10 μg/mL of the phthalate mixture (PM10), 0.1 μg/mL of the metabolite mixture (MM0.1), or 10 μg/mL of the metabolite mixture (MM10) for 96 hours. At the end of the culture period, RNA from the cells was subjected to bulk RNA-seq (A) Heatmap of differential expression of genes with global FDR p-value < 0.2 (B) Genes with global < 0.2 are highlighted in red (up-regulated) or blue (down-regulated) for each treatment group compared to the control (C) Number of differentially expressed genes for each treatment group compared to the control when the global FDR p-value was restricted to < 0.05 for further analysis (D) Venn diagram of overlapping genes. Abbreviations: DMSO, dimethylsulfoxide; FDR, false discovery rate; PM, phthalate mixture; MM, metabolite mixture

We used DAVID for functional annotation of differentially expressed genes. As mixtures of phthalate diesters and monoesters have been previously shown to exert a variety of effects on the ovary, sometimes the same and sometimes different (Meling *et al*., 2020; Warner *et al*., 2021; Zhou and Flaws, 2017), we analyzed the differentially expressed gene lists for the phthalate treatments separately (each treatment group, **Table S3**) and together (all 407 genes identified using ANOVA, **Table S2**). We did not further analyze the 0.1 ug/mL PM treatment group due to lack of differentially expressed genes.

Analysis of gene ontology (GO) enriched functions and Kyoto Encyclopedia of Genes and Genomes (KEGG) pathways for all 407 genes and the individual 10 μg/mL PM and 10 μg/mL MM treatment groups compared to control with adjusted p-values (Benjamini) of < 0.05 are shown in **Table 1**. Ten pathways and/or terms were enriched for the full gene list, including lipid metabolic process (FDR = 7.74E-08), PPAR signaling (FDR = 5.12E-05), and positive regulation of smooth muscle cell proliferation (FDR = 0.006). Other pathways involved lipids and/or metabolism, including cholesterol metabolism (FDR = 0.019), glycerophospholipid metabolism (FDR = 0.031), and metabolic pathways (FDR = 0.04). Three and four pathways and/or terms were enriched for 10 μg/mL PM and 10 μg/mL MM treatment groups compared to control. Lipid metabolic process, PPAR signaling, and fatty acid metabolic process were enriched for both treatments (**Table 1**), and cell adhesion (FDR = 0.0056) was enriched only for PM 10.

**Table 1:** Analysis of biological process gene ontology (GO) and Kyoto Encyclopedia of Genes and Genomes (KEGG) pathways of 407 differentially expressed genes.

| Category | ID | description | Count | p Value | Benjamini |
| --- | --- | --- | --- | --- | --- |
| *All treatment groups: 407 genes* |  |  |  |  |  |
| GOTERM_BP_DIRECT | GO:0006629 | **lipid metabolic process** | 44 | 2.77E-11 | 7.74E-08 |
| KEGG_PATHWAY | mmu03320 | **PPAR signaling pathway** | 14 | 1.98E-07 | 5.12E-05 |
| GOTERM_BP_DIRECT | GO:0048661 | positive regulation of smooth muscle cell proliferation | 11 | 4.39E-06 | 0.00613102 |
| GOTERM_BP_DIRECT | GO:0009410 | response to xenobiotic stimulus | 20 | 1.09E-05 | 0.01011471 |
| KEGG_PATHWAY | mmu04979 | Cholesterol metabolism | 8 | 1.86E-04 | 0.01898861 |
| KEGG_PATHWAY | mmu04974 | Protein digestion and absorption | 11 | 2.81E-04 | 0.01898861 |
| KEGG_PATHWAY | mmu04512 | ECM-receptor interaction | 10 | 2.93E-04 | 0.01898861 |
| KEGG_PATHWAY | mmu00564 | Glycerophospholipid metabolism | 10 | 6.01E-04 | 0.0311495 |
| GOTERM_BP_DIRECT | GO:0042127 | regulation of cell proliferation | 15 | 4.55E-05 | 0.03171773 |
| KEGG_PATHWAY | mmu01100 | Metabolic pathways | 59 | 0.00109514 | 0.04727357 |
| *10 ug/mL PM vs control: 123 genes* |  |  |  |  |  |
| GOTERM_BP_DIRECT | GO:0006629 | **lipid metabolic process** | 20 | 2.71E-08 | 3.13E-05 |
| KEGG_PATHWAY | mmu03320 | **PPAR signaling pathway** | 8 | 8.99E-06 | 0.00140209 |
| GOTERM_BP_DIRECT | GO:000663 | fatty acid metabolic process | 9 | 2.07E-05 | 0.01193019 |
| *10 ug/mL MM vs control: 90 genes* |  |  |  |  |  |
| KEGG_PATHWAY | mmu03320 | **PPAR signaling pathway** | 8 | 2.14E-07 | 2.44E-05 |
| GOTERM_BP_DIRECT | GO:0007155 | cell adhesion | 13 | 7.43E-06 | 0.00561454 |
| GOTERM_BP_DIRECT | GO:000663 | fatty acid metabolic process | 8 | 1.97E-05 | 0.00744707 |
| GOTERM_BP_DIRECT | GO:0006629 | **lipid metabolic process** | 13 | 3.96E-05 | 0.00996771 |

The 0.1 μg/mL MM treatment group had 16 differentially expressed genes, which we also subjected to pathways analysis to see if any of the same processes were identified as the 10 ug/mL treatment groups. Multiple terms related to transcription regulation were identified (**Table S4**), but none overlapped with the other treatment groups or appeared in the analysis from all 407 genes. Due to the small gene list, we did not further analyze this treatment group on its own.

Finally, we used the function annotation clustering feature in DAVID to group similar annotations together for a different look at the differentially expressed genes lists. For this analysis, we expanded the annotation analysis to include UniProt, Biocarta, Interpro, and other DAVID-default functional databases for a broader look at pathways and interactions. Notably, the most enriched cluster (enrichment score 5.01) for the full differentially expressed gene list included terms related to endoplasmic reticulum (**Table S5**), followed by extra cellular space (enrichment score 3.32). Performing the same analysis on each 10 μg/mL gene list revealed PPAR in the most enriched cluster for the PM treatment (enrichment score 3.45) and endoplasmic reticulum as the second most enriched cluster (enrichment score 3.42), whereas the MM treatment’s most enriched clusters are related to extracellular matrix (enrichment score 2.73 and 2.54).

### PPAR analysis

The related terms of PPAR signaling and lipid metabolic processes appeared repeated across gene function analyses, so we chose to focus on these for further study. These are related processes with multiple shared signaling factors. We used the ppargene.org database to identify verified and predicted PPAR target genes. Of the 15 total genes identified by DAVID as part of the PPAR signaling pathway, 12 are verified or predicted to be transcriptionally regulated by PPAR alpha, delta, or gamma (**Table S6**). All of the 8 genes in the PPAR pathway from the 10 μg/mL PM treatment group were upregulated (*Plin2, Plin1, Acsbg1, Hmgcs1, Cd36, Pltp, Angptl4,* and Plin5). Of the 8 genes in the PPAR pathway from the 10 μg/mL MM treatment group, 7 were upregulated (*Plin2, Plin1, Cd36, Scd4, Angptl4, Aqp,7* and *Cpt1b*) and *Me3* was downregulated, and four genes were commonly upregulated between the two treatment groups (*Plin2, Plin1, Cd36,* and *Angptl4*).

Of the 45 total genes from the lipid metabolic process GO term, 29 (64%) are verified or predicted PPAR targets (**Table S7**). All 13 genes in the lipid metabolic process term from the 10 μg/mL MM treatment group, all were upregulated, whereas 17 of 20 were upregulated from the 10 μg/mL PM treatment group. Seven genes (*Acsf2, Angptl4, Hadha, Hadhb, Plin1, Mgll*, and *Plpp3*) were upregulated in both treatment groups.

To further explore the transcriptional regulators of the differentially expressed genes, we analyzed the differentially expressed gene lists from 10 μg/mL PM, 10 μg/mL MM, and all 407 genes using the epigenetic Landscape In Silico deletion Analysis (LISA) tool at cistrome.org (Qin *et al*., 2020). For the full gene list, the top regulator was CEBPB (p = 3.93E-19), a PPARG cofactor (**Table S8**). PPARG and the PPAR heterodimerization partner RXRA were also identified as regulators (p = 2.40E-15 and p = 4.51E-12, respectively). PPARG was the top transcription factor for both the PM10 and MM10 treatment groups (p = 5.16E-15 and 6.16E-13, respectively), with the PPAR binding protein MED1, PPARA, RXRG, and CEPBP (p = 6.90E-11, 2.25E-08, 1.59E-07, 1.22E-06, respectively) identified as common transcription factors in the PM10 treatment group. RXRA, MED1, CEPBP, and CEBPA (p = 1.98E-10, 5.62E-09, 4.80E-07, 6.42E-07, respectively) are identified as common transcription factors for MM10. NR3C1, the glucocorticoid receptor, also appears in all three lists (all groups: p = 9.86E-18; PM10: p = 1.12E-05; MM10: p = 5.51E-10), as well as the ovarian transcription factor FOXL2 (all groups: p = 1.63E-15; PM10: p = 3.23E-07; MM10: p = 2.14E-05).

### qPCR confirmation of RNA-seq results

We selected a variety of differently expressed genes (*Cd36, Angptl4, Pdk4, Plin1, Plin2, Mgll, Hp, Lpl,* and *Pltp*) for qPCR analysis to verify the RNA-seq analysis data (**Figure 7**). Overall, the RNA-seq data are consistent with the qPCR analysis of these genes. The qPCR analysis includes all treatment groups from the culture experiments and reveals statistically significant increases in the 100 μg/mL treatment groups for the phthalate mixture (*Cd36*, *Angptl4*, *Pdk4*, *Plin1*, *Mgll*, *Hp*, and *Pltp*) and metabolite mixture (*Plin1*, *Mgll*, *Hp*, and *Pltp*). Fewer genes were impacted at the 100 μg/mL dose by the metabolite mixture compared to the phthalate mixture, but *Plin1* and *MgII* had noticeably larger fold change differences in due to exposure to 100 μg/mL MM compared to 100 μg/mL PM. For *Angptl4*, *Hp*, and *Lpl*, statistically significant increases in expression were observed at the 1 μg/mL MM dose compared to control.

**Figure 7:**
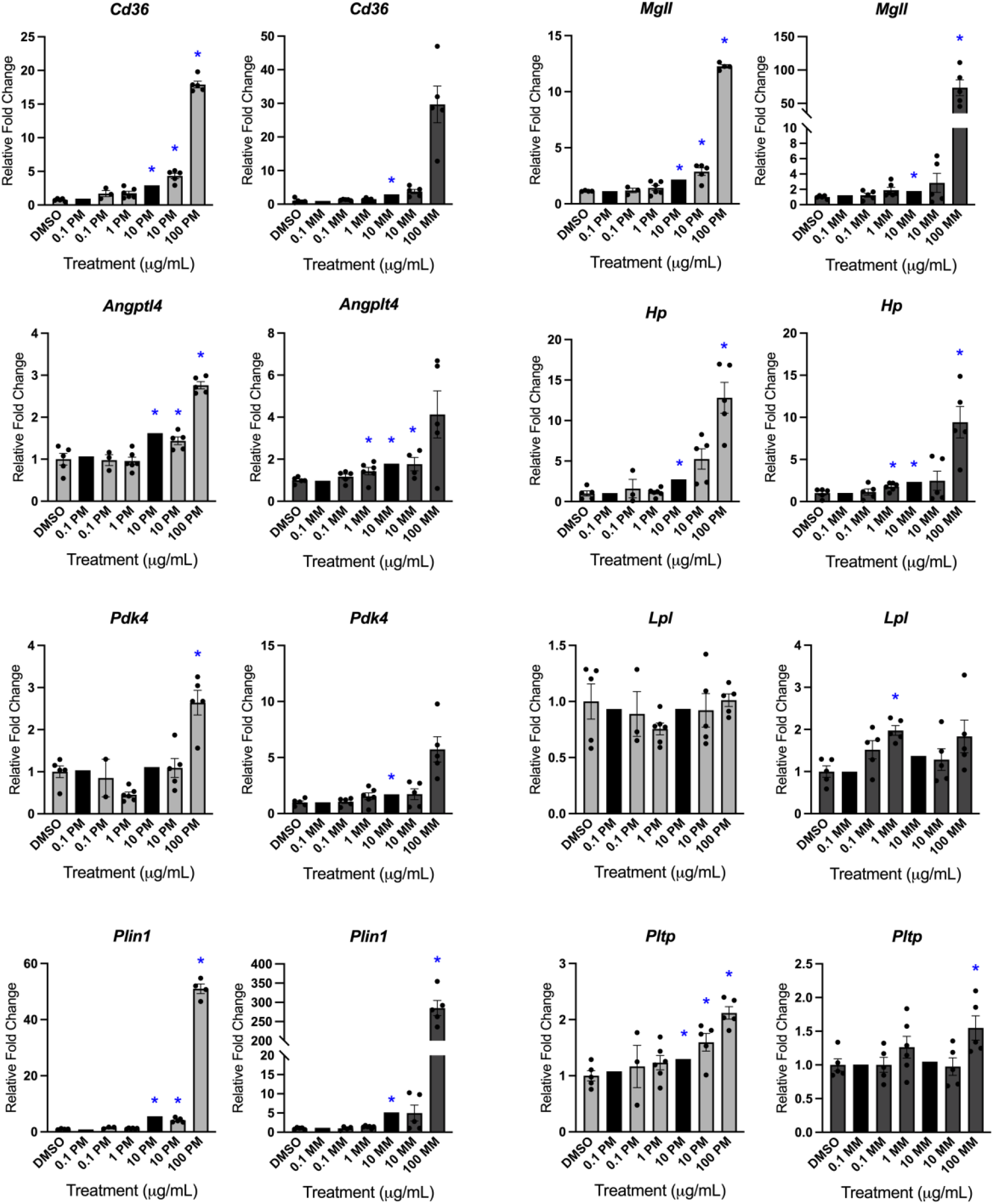
Expression of select differentially expressed genes from the RNA-seq analysis were confirmed using qPCR. Granulosa cells were harvested from adult female mice and cultured for 96 hours in the presence of the phthalate mixture (PM) or metabolite mixtures (MM) at 0.1 – 100 μg/mL. RNA extracted from the cultured granulosa cells were subjected to qPCR (gray bars) or RNA-sequencing (black bars). Graphs represent mean ± SEM from 3–6 separate experiments. Relative fold changes of each gene normalized to Rn18s are shown for qPCR. Fold change from RNA-seq is reported as an average of all samples (see Table S2). Asterisks (∗) indicate significant differences from the control (p ≤ 0.05).

## Discussion

In this paper, we report on our studies focusing on the granulosa cell, in which we find that granulosa cells are largely not contributing to phthalate metabolism, but are targets of phthalate toxicity through lipid metabolism-related pathways.

This study directly follows up our previous work on the metabolism of phthalates in the ovary (Warner *et al*., 2019). Previously, we found that whole antral follicles expressed two enzymes that are possible biotransformers of phthalates, ALDH1A1 and LPL. As granulosa cells are the most abundant cell type in the follicle and surround the antrum, in which phthalates have been measured, we hypothesized that these enzymes were expressed in granulosa cells. To test this, we performed immunohistochemistry on whole, uncultured young adult mouse ovaries. Surprisingly, ALDH1A1 was strongly expressed in the stroma surrounding the follicles and in the theca cells of antral follicles. This enzyme was not observed in neonatal ovaries, which were stained to analyze primary and primordial follicle populations, nor was it observed in granulosa cells of antral follicles. Theca cells derive from stroma cells and become more numerous and differentiated at the antral follicle stage, so it is unsurprising that expression is shared between the two cell types. On the other hand, LPL was observed in the stroma and granulosa cells of primordial and preantral follicles, but faded away at the antral stage. The lack of expression of either enzyme in granulosa cells from antral follicles in consistent with our observation that granulosa cells in culture are largely not transforming parent phthalates into metabolites. The fact that most metabolism of phthalates in the ovary is likely occurring in the stroma, rather than the follicles themselves, suggests that our previous analysis of phthalate metabolism in cultured antral follicles grossly underestimated the biotransformation capacity of adult ovaries. The interesting observation of lack of metabolism by mature granulosa cells may provide a protective effect for the oocyte as it approaches ovulation. The strong expression of metabolizing enzymes in the stroma of adult ovaries suggests that the ovary may be attempting to detoxify xenobiotics arriving via the bloodstream before they reach the sensitive follicles.

Although literature reports implicate LPL and ALDH1A1 as metabolizers of phthalates (Albro and Thomas, 1973; Ito *et al*., 2005) and our measurements of these enzymes are consistent with our calculations of phthalate metabolism, we have not confirmed that these are the predominant enzymes performing phthalate metabolism in the ovary. We attempted to culture antral follicles in the presence of a lipase inhibiter, the surfactant poloxamer-407 (Johnston, 2010; Johnston and Goldberg, 2006; Johnston and Palmer, 1993), but lipases perform essential processes in the ovary; overall follicle growth was too inhibited to draw conclusions about the role of lipases in phthalate metabolism (data not shown.)

This study finds that granulosa cells are largely not transforming parent phthalates into metabolites, with less than 20% of phthalates converted into monoester metabolites in culture across the 1, 10, and 100 μg/mL PM treatment groups, compared to up to 85% conversion in neonatal ovary culture and antral follicle culture (**Figure 8**). This study does not report metabolism in the 0.1 μg/mL treatment group because of contamination identified by the mass spectrometry analysis in the vehicle control group at similar concentrations as the phthalate measurements in the 0.1 μg/mL PM treatment group. Unfortunately, we do not know if the contamination (DEP, DEHP, and DiNP) was introduced during the culture process or during the mass spectrometry analysis. Therefore, although gene expression results from the 0.1 μg/mL treatment groups are reported in this study, we do not know if there are few statistically significant effects at this dose because of a lack of toxicity or because the dose is indistinguishable from background contamination in the vehicle control.

**Figure 8:**
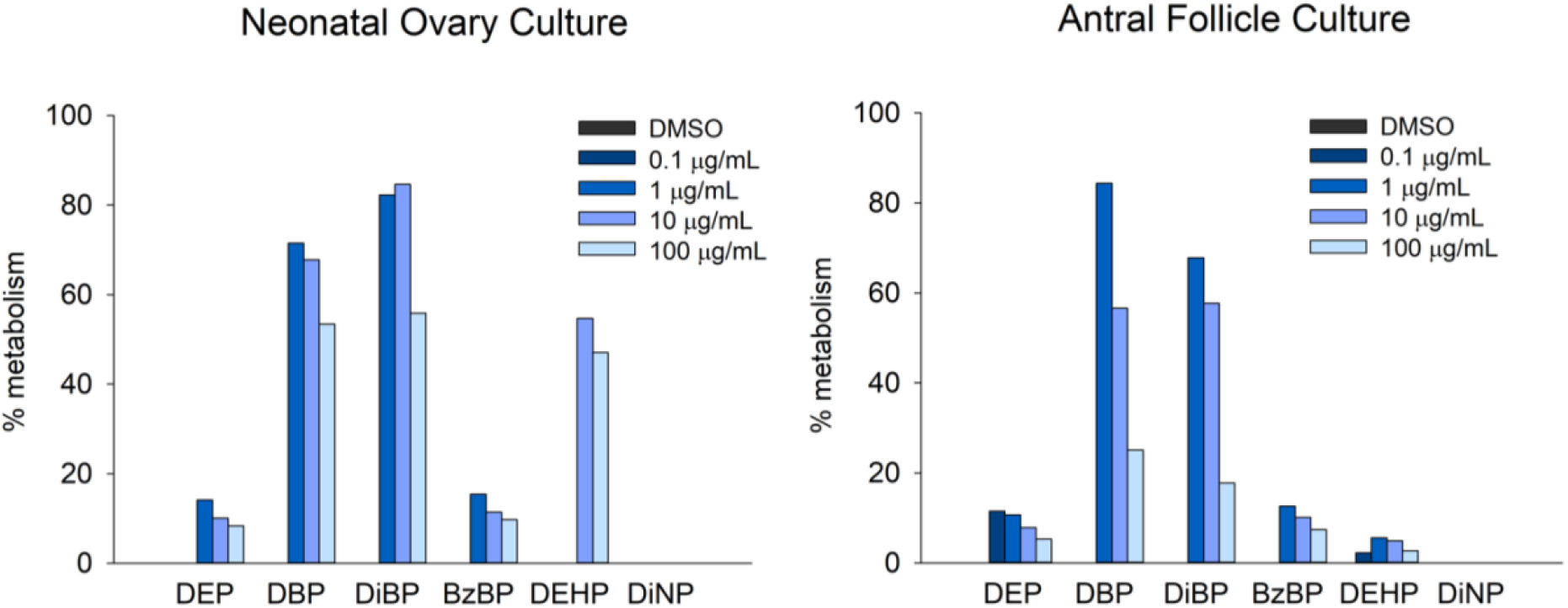
Summary figure of the percent metabolism of individual phthalates measured by LC-MS/MS following culture of whole postnatal day 8 ovaries (left) or adult mouse antral follicles (right) in the presence of the phthalate mixture (0.1–100 μg/mL) for 96 hrs are reported in Warner *et al.,* 2019.

It is widely reported that phthalates are bioactivated into monoester metabolites that are more toxic than the parent diesters, but given the metabolic capacity of the ovary and the ease with which phthalates are hydrolyzed, it is still unknown whether diester phthalates cause direct ovarian toxicity. This study, which identified granulosa cells from antral follicles as one of the ovarian cell types least able to bioactive phthalates, provides insight into this question. We performed global gene expression analysis on RNA extraction from the 0.1 and 10 μg/mL phthalate mixture and metabolite mixture treatment groups. Few differentially expressed genes were observed for the 0.1 μg/mL treatment groups, but 123 and 90 differentially expressed genes were identified for the 10 μg/mL PM and MM treatment groups, respectively. We confirmed the RNA-seq gene expression data by performing qPCR on 8 selected differentially expressed genes. Overall, there were a similar number of statistically significantly affected treatment groups across both mixtures, although the impacted doses and treatments were not always the same. The phthalate mixture, but not the metabolite mixture, caused statistically significant increases in gene expression of *Cd36*, *Angptl4*, and *Pdk4* at 100 μg/mL and *Pltp* at 10 μg/mL. In converse, the metabolite mixture, but not the phthalate mixture, caused statistically significant increases in *Angptl4, Lpl,* and *Hp* at 1 μg/mL and *Pdk4* at 10 μg/mL. If the biotransformation products of the PM doses were solely responsible for observed toxicity, we would expect to see effects from the PM at 10-100x higher doses than is observed from MM treatment. We see this trend, for example, with *Angptl4*, *Pdkr4*, and *Hp*, where lower doses of the MM cause statistically significant effects compared to the PM, but this relationship does not hold true for all genes subjected to qPCR. In addition, we observed more statistically significant differentially expressed genes from the 10 μg/mL PM treatment than the 10 μg/mL MM treatment in the RNA-seq analysis. Similarly, more statistically significant disruptions to gene expression of steroidogenic enzymes were observed from PM treatment than MM treatment. Overall, the robust effects from the PM that are different from the MM in this model system in which metabolism is limited compared to whole tissues suggest that diesters are not neutral parties; the complex mixture of the PM, including both the parent phthalates and a small fraction of bioactivated metabolites, are contributing to phthalate toxicity differently from metabolites alone.

Previous studies have identified PPAR signaling as a target of phthalate toxicity and potential mechanism of action in the ovary, particularly in granulosa cells (Lovekamp-Swan *et al*., 2003; Meling *et al*., 2022). In silico docking studies predict that phthalates bind to PPAR subtypes, as well as their dimerization partners RXRs (Meling *et al*., 2022; Sarath Josh *et al*., 2014). In Meling *et al.,* 2022, we previously showed that individual high molecular weight phthalate monoesters can significantly induce the expression of PPAR target genes *Fabp4* and *Cd36* at 40 and 400 μM (approximately 10 and 100 μg/mL, respectively) in cultured mouse granulosa cells. In the present study, *Cd36* was the most significantly differentially expressed gene overall and by the 10 μg/mL PM and MM mixtures. Only MEHP treatment alone at 40 μM (10 μg/mL) significantly increased the expression of *Cd36* compared to control in Meling et al., whereas in this study both of the mixtures at 10 μg/mL significantly increased *Cd36* expression at the same total dose, showing that the effects of the components of the mixture is more than additive; the mixture components are synergistically increasing toxicity compared to individual phthalate treatments. Interestingly, only monoester metabolites (notably, MEHP) have been shown to be able to interact with PPAR ligand binding domains, not diesters, suggesting that either the monoester are active at very low concentrations in the PM, or the diesters are acting through other mechanisms (Desvergne *et al*., 2009).

Pathway analysis of the differentially expressed genes lists from RNA sequencing identified PPAR signaling and lipid metabolism as impacted pathways for all gene lists analyzed (all treatment groups, 10 μg/mL PM, or 10 μg/mL MM treatment each vs control, **Table 1**). We chose to analyze each of these gene lists to identify common effects from phthalates and effects that are unique to each mixture. PPAR signaling and lipid metabolism are closely related; PPAR receptors’ endogenous ligands are fatty acids, and they act as transcription factors, as PPAR-RXR heterodimers, for fatty acid synthesis and metabolism, lipid storage, and adipogenesis (Desvergne *et al*., 2009). We identified many differentially expressed genes that are PPAR targets (**Tables S6** and **S7**), and further confirmed PPAR and RXR isoforms and their cofactors as major transcription factors regulating the differentially expressed gene lists. Overall, our results of this non-targeted RNA-sequencing experiment are consistent with the extensive body of literature implicating PPAR signaling as a mechanism of phthalate action.

It is worth considering what implications the interactions of phthalates with PPARs and lipid metabolism have for the metabolism of phthalates in the ovary, given that one of our key enzymes of interest, LPL, has an important role in lipid metabolism by catalyzing the hydrolysis of triglycerides into fatty acids. The interaction of phthalates with PPAR receptors and resulting dysregulation of downstream target genes may serve to reduce the ability of the ovary to metabolize phthalates. Notably, the second-most statistically significantly impacted gene across all three gene lists (after *Cd36*) was *Angptl4*. ANGPTL4 is an inhibitor of LPL (Wu *et al*., 2021). *Lpl* was statistically significantly upregulated following 1 μg/mL MM treatment, but not 10 or 100 μg/mL, compared to control, whereas *Angptl4* expression continued to increase with increasing dose. *Angptl4* induction may be preventing the ovary from increasing expression of metabolizing enzymes.

We additionally identified endoplasmic reticulum and extracellular matrix terms in our functional annotation of differentially expressed genes from all three gene lists. The mechanisms through which phthalates may interact with genes clustered in these terms are less well understood compared to PPARs and lipid metabolism, and in fact may be related to PPAR mechanisms of action, as many of the genes are PPAR target genes. Endoplasmic reticulum stress can lead to dysregulation of lipid metabolism (Basseri and Austin, 2011). Induction of PPARs is associated with prevention and/or reduction of endoplasmic reticulum stress (Ikeda *et al*., 2015; Salvadó *et al*., 2014; Soliman *et al*., 2019). From our data set, it is not clear if the association between endoplasmic reticulum associated genes and pathways and phthalate treatment is related to endoplasmic stress or other dysregulation of endoplasmic reticulum function; key markers of endoplasmic reticulum stress in the ovary, including heat shock 70 kDa protein 5 (*Hspa5*), activating transcription factor 4 (*Atf4*), *Atf6*, and C/EBP homologous protein (*Chop*) were not differentially expressed in the RNA-seq results (Takahashi *et al*., 2017). Phthalates, particularly DEHP/MEHP and DBP/MBP, have been associated with endoplasmic reticulum stress in cell lines and testes (Ozkemahli *et al*., 2023; Pan *et al*., 2019; Peropadre *et al*., 2015; Yöntem *et al*., 2024; Zhang *et al*., 2016, 2019; Zhu *et al*., 2021). Future studies are necessary to investigate whether endoplasmic reticulum stress may play a role in phthalate toxicity in the ovary.

In conclusion, this study confirms, through non-targeted RNA-sequencing analysis of mouse ovarian granulosa cells exposed to phthalate mixtures, that PPAR signaling and lipid metabolism are targets of phthalate toxicity. Our phthalate mixtures exerted different effects, emphasizing the importance of considering complex mixtures when assessing the toxicity of phthalates. Overall, granulosa cells are not significantly contributing to phthalate metabolism compared to other cell types in the ovary, but are targets of phthalate toxicity. Future studies will investigate the molecular mechanisms through which phthalates interfere with PPAR signaling and their interactions with other cell types in the ovary to help provide mechanistic information on phthalate toxicity in the ovary for use in adverse outcome pathways and risk assessment.

## Data links

Raw RNA-sequencing files are available through SRA at BioProject: PRJNA1096659

All raw immunohistochemistry images from the study are available at Open Science Framework (https://osf.io/8r3aw/?view_only=b60aa9b030294e6fb50337b7361967d5)

## Acknowledgements

Thank you to the members of the Flaws Lab at the University of Illinois at Urbana-Champaign, especially Drs. Daryl Meling and Mary Laws, Dr. Jenny Drnevich from the High Performance Computing in Biology (HPCBio) and The Roy J. Carver Biotechnology Center at the University of Illinois at Urbana-Champaign, and all members of the EDC Lab at NJIT.

